# Transcriptome profiles of telocytes along the intestinal crypt-villus axis

**DOI:** 10.64898/2025.12.03.692064

**Authors:** Amal Gharbi, Noa Corem, Sharona Elgavish, Inbar Plachkes, Abrar Jamous, Marco Canella, Moriya Shmuel, Hadar Horowitz, Jianmei Tan, Hongqian Chen, Bing Su, Mahmud Abu-Gazala, Noam Shussman, Michal Shoshkes-Carmel

## Abstract

**Background & aims:** Intestinal epithelial cells rely on a complex array of signals from the stromal microenvironment to determine their fate and function along the crypt-villus axis, but the precise cellular sources and combinations of signals regulating each position remain poorly defined.

**Methods:** Here, we present a high-resolution atlas of a unique mesenchymal cell type previously identified as telocytes, mapped along the crypt-villus axis using single-cell RNA sequencing integrated with *in-situ* approaches and reporter mouse models.

**Results:** We identify four spatially distinct telocyte subtypes: crypt, villus base, villus mid and villus tip, each with a unique gene expression profile. Crypt telocytes are enriched for canonical Wnt signaling components that support stem and progenitor cell maintenance, while villus base telocytes are a source of Bmps (bone morphogenic proteins) and express contractility-associated genes, suggesting a role in tissue architecture. Villus mid telocytes display signatures of immune regulation and inflammation, while villus tip telocytes are specialized for the regulation of ribonucleoprotein complexes and nutrient sensing.

**Conclusions:** This atlas provides a foundational resource for understanding how diverse telocyte populations coordinate signaling environments and shape intestinal homeostasis through both epithelial interactions and extensive crosstalk among themselves.

## Introduction

The small intestine is lined by a single layer of epithelial cells organized into repeating crypt-villus units. This structured arrangement is crucial for protecting stem and progenitor cells in the crypt from the harsh conditions of the intestinal lumen, maximizing nutrient absorption, and maintaining a robust, continuous barrier against physical stress and invading pathogens.

Along the crypt-villus axis, specific cell types and functions are restricted to defined locations. Remarkably, this intricate pattern is preserved despite the intestine’s rapid and continuous renewal. Stem and progenitor cells at the crypt divide continuously to produce new cells, which then migrate upward along the villus and are eventually shed at the tip. As a result, the epithelial layer is in perpetual motion. Yet, throughout this dynamic process, the overall spatial organization remains stable. This stability is achieved because stromal signals along the crypt-villus axis repeatedly instruct epithelial cells to adopt specific roles as they migrate toward the villus tip.

Among intestinal stromal cells, telocytes have emerged as a distinct mesenchymal population, notable for their exceptionally long cellular projections known as telopodes^1–6^. Intestinal telocytes, which express the transcription factor Foxl1, form an interconnected subepithelial network that plays an essential role in both maintaining the stem cell pool and directing the differentiation of enterocytes^6–10^. Telocytes are recognized for compartmentalizing signaling molecules along the crypt-villus axis^7,8^. However, their molecular diversity necessitates single-cell resolution for full characterization. Despite advances in characterizing stromal cells^8,11–15^, the low expression of Foxl1 and the molecular diversity of telocytes have limited their comprehensive analysis using conventional single-cell approaches. To overcome these challenges, here we employed a Foxl1Cre-driven reporter mouse model combined with single cell RNAseq and single molecule RNA fluorescence in-situ hybridization (smFISH) to enrich, visualize and molecularly profile telocyte populations *in situ*. We report distinct telocyte subtypes with defined expression patterns along the crypt-villus axis, explaining how telocytes can contribute to determining intestinal epithelial cell fate.

## Results

### Foxl1-expressing telocytes form a subepithelial network that extends from the crypt base to the villus tip in the small intestine

While previous studies established that PDGFRα+ telocytes creates a continuous subepithelial network in mice^7,8^, it remained unclear whether this organization is conserved in the human intestine. To address this, we analyzed fresh tissue samples from normal margines of colectomies and intestinal resections, examining both large and small intestines following clearing and PDGFRα staining (**Figure 1A and B**). In the colon, PDGFRα+ telocytes formed a continuous network closely associated with crypts (**Figure 1A**), while in the small intestine, the network spanned the entire crypt-villus axis (**Figure 1B**). This distribution mirrored the mouse pattern, suggesting a conserved role across species. Beyond proximity to the epithelium, telocytes also contacted CD45+ immune cells (**Figure 1A**) and αSMA+ myofibroblasts in the crypt region, while myofibroblasts in the villus localized to the core (**Figure 1B**).

**Figure 1.**
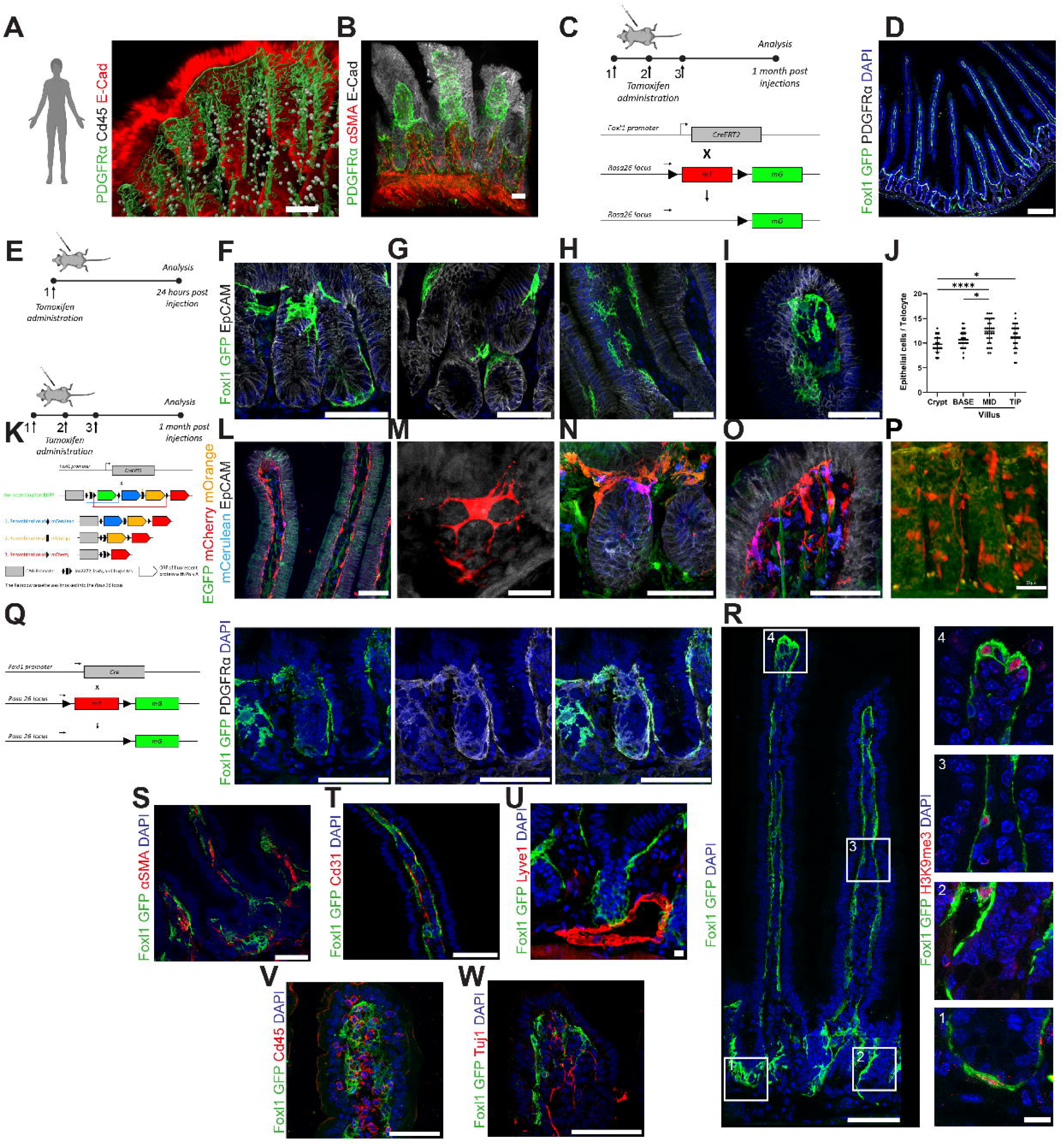
Foxl1-expressing telocytes form a subepithelial network that extends from the crypt base to the villus tip in the small intestine. (A-B) Whole-mount immunofluorescence imaging of cleared human colon (A) and small intestine (B). Panel A depicts a 3D volume reconstruction generated in IMARIS, while Panel B shows a z-stack projection. PDGFRα expression highlights a subepithelial network along the crypts in the colon and the crypt-villus axis of the intestine. E-Cadherin marks the epithelial layer, CD45 labels immune cells, and αSMA marks myofibroblasts, demonstrating close spatial relationships between subepithelial PDGFRα+ telocytes, epithelial cells, immune cells, and myofibroblasts. (C) Schematic illustrating the experimental design: tamoxifen induction in Foxl1-CreERT2; Rosa-mTmG mice drives membrane-bound GFP expression in Foxl1+ cells. (D) Immunofluorescence of Foxl1CreERT2; Rosa-mTmG mouse jejunal sections shows GFP+ telocytes forming a subepithelial network along the crypt-villus axis. (E) Schematic showing that single tamoxifen induction labels Foxl1+ telocytes inefficiently in Foxl1CreERT2; Rosa-mTmG mice. (F-I) Immunofluorescence for GFP in Foxl1CreERT2; Rosa-mTmG jejunal sections following single tamoxifen induction reveals individual telocytes along the crypt-villus axis. (J) Quantification of epithelial cell contacts by a single telocyte at defined locations along the crypt-villus axis, measured in 30 μm-thick sections under inefficient Foxl1CreERT2; Rosa-mTmG and efficient Foxl1CreERT2; Rosa-Rainbow tamoxifen induction (n=3/group, <10 axes/bar; ****P<0.0001, unpaired two-tailed t-test). (K) Schematic depicting tamoxifen-induced expression of membrane-bound multispectral Rainbow fluorescence in Foxl1+ cells (Foxl1CreERT2; Rosa-Rainbow mice). (L-O) Immunofluorescence of Foxl1CreERT2; Rosa-Rainbow jejunal sections stained for RFP, single telocytes are visualized along the villus (L), crypt (M-N), and villus tip (O). (P) Whole-mount immunofluorescence of Foxl1CreERT2; Rosa-Rainbow colon stained for RFP highlights single telocytes along colon crypts. (Q) Schematic showing how membrane-bound GFP is expressed in Foxl1+ cells (Foxl1Cre; Rosa-mTmG). Immunofluorescence for GFP demonstrates limited reporter signal in crypts, whereas PDGFRα staining captures the full extent of telocyte processes. (R) Immunofluorescence for GFP and H3K9me3 in Foxl1Cre; Rosa-mTmG jejunal sections demonstrates relatively high heterochromatin content in Foxl1-GFP nuclei along the crypt-villus axis. (S-W) High-resolution imaging reveals GFP+ telocytes in close proximity to myofibroblasts (αSMA, S), blood vessels (Cd31, T), lymphatic vessels (Lyve1, U), immune cells (CD45, V), and neurons (Tuj1, W). Scale bars: 50 μm.

To further characterize the telocyte network in mice, we employed the inducible Foxl1CreERT2 mouse model^16^ crossed with the Rosa-mTmG reporter. Administration of three tamoxifen injections produced membrane-tagged GFP labeling across the crypt-villus subepithelial network (**Figure 1C and D**), in contrast to the sparse recombination observed in the same model crossed with the Rosa-YFP reporter^17^. This highlights the utility of membrane-tagged reporters for visualizing telocyte structure. Notably, GFP signal was consistently present throughout the villus, while labeling in the crypt region was distinctly sparse. To resolve individual telocyte architecture, we applied a single, inefficient tamoxifen induction (**Figure 1E-J, Video1 and 2**) or used Foxl1CreERT2; Rosa-Rainbow mice with efficient induction, producing multispectral labeling (**Figure 1K-P, Video 3**). Individual telocytes were arranged in a thin layer at the epithelial base along the crypt-villus axis, supporting previous findings. Villus telocytes displayed a stellate morphology with multiple branched extensions forming a continuous network (**Video 2 and 3**), while crypt telocytes appeared flat and thin with extremely slender, short processes (**Figure 1M and Video 1**), consistent with descriptions of a “pericryptal mesenchymal syncytium”^18^. At the villus tip, telocytes adopted an orthogonal orientation relative to the epithelium (**Figure 1I and O**). Each telocyte interacted with an average of ten epithelial cells, based on quantification from the jejunum (**Figure 1J**, see Methods), though accurate enumeration is challenging due to their fine, oriented processes. These findings suggest that telocytes might play a role in arranging the crypt-villus axis by dividing the epithelium into segments. This organization could help direct epithelial cell behavior at specific locations in the intestine. However, further studies are needed to test this intriguing possibility.

Similar structural features were detected in mouse colon (**Figure 1P**), with telocytes exhibiting small, flat cell bodies or long processes extending up to a third of a crypt’s length. The sparse reporter signal observed in the crypt region reflects both the flat morphology of crypt telocytes and a technical limitation of the Rosa-mTmG/Rainbow reporter, which restricts fluorescence to the inner membrane and thus fails to highlight thin, flat structures that PDGFRα immunostaining can capture more completely (**Figure 1Q**). Therefore, the apparent paucity of reporter signal does not indicate telocyte absence, but results from the distinct crypt telocyte architecture and fluorophore localization constraints. Consistent with electron microscopy studies, telocyte nuclei exhibited relatively high heterochromatin content as determined by H3K9me3 staining (**Figure 1R**).

Telocytes also maintained close associations with aSMA+ myofibroblasts, CD31+ vascular vessels, Lyve1+ lymphatic vessels, CD45+ immune cells, and Tuj1+ neurons (**Figure 1S-W**). These contacts underscore their potential role in mediating epithelial, stromal, vascular, immune, and neuronal interactions. Collectively, our structural analysis reveals an intricate subepithelial network formed by telocytes, supporting their role as coordinators of complex intestinal signaling.

### Single-cell transcriptomics distinguishes four telocyte subtypes

To assess whether Foxl1, a transcription factor important for telocyte function and a recognized telocyte marker^19–22^, is uniformly expressed along the crypt-villus axis, we used a dual approach: Foxl1Cre^23^; Rosa-mTmG reporter mice and single molecule RNA fluorescence in situ hybridization (smFISH) to quantify Foxl1 transcripts (**Figure 2A**). This analysis revealed that villus telocytes possess higher Foxl1 transcript abundance than those in the crypt (**Figure 2B**).

**Figure 2.**
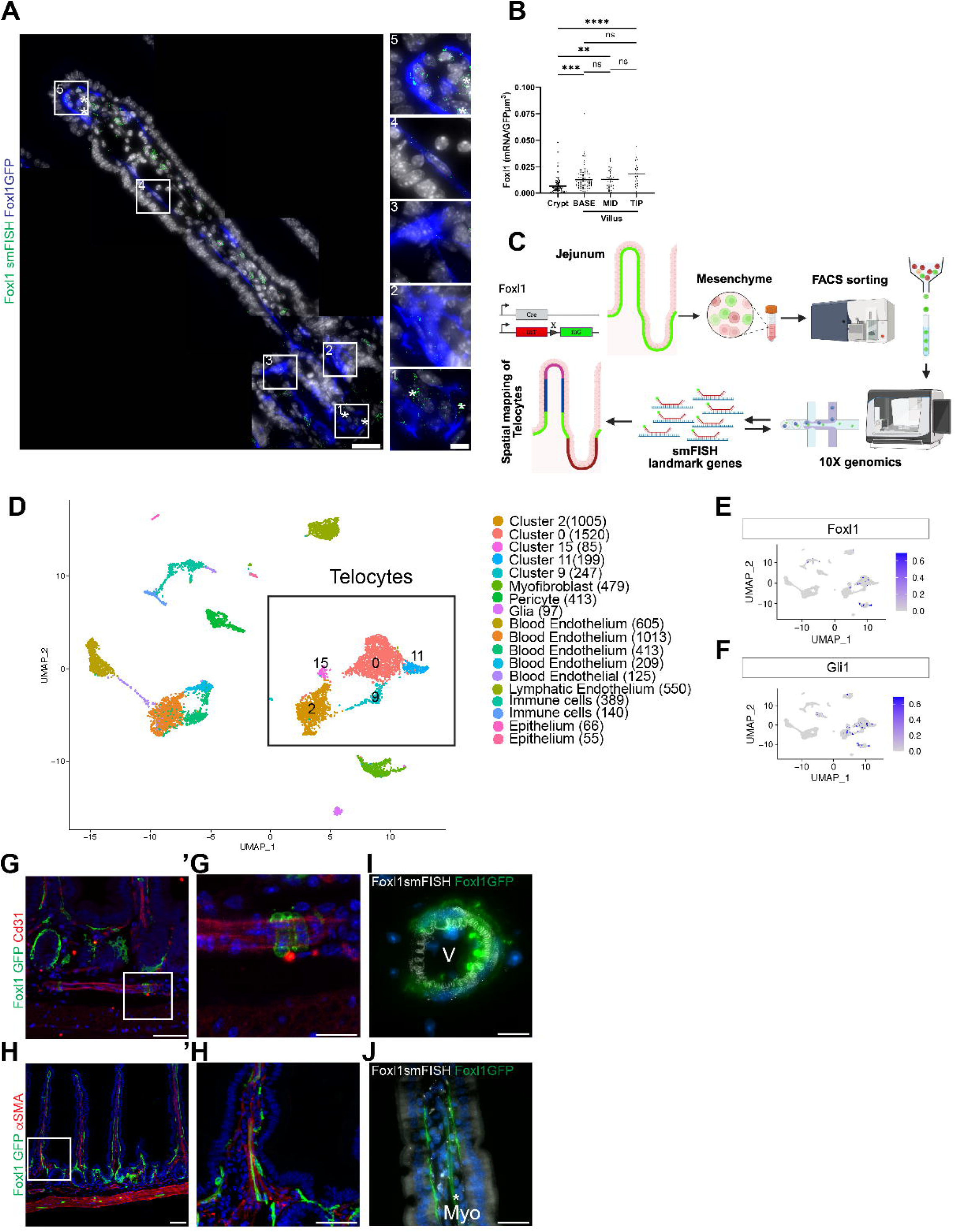
Single-cell transcriptomics distinguishes four telocyte subtypes. (A) smFISH imaging of mouse jejunum sections from Foxl1Cre; Rosa-mTmG mice, hybridized for Foxl1 (green dots), demonstrates Foxl1 mRNA expression in GFP+ telocytes (blue) positioned at the crypt, villus base, villus mid, and villus tip. (B) Quantification of Foxl1 mRNA concentrations in telocytes along the crypt-villus axis by smFISH, revealing spatial variation (n=3 mice/group, <10 crypt-villus axes/bar, ****P<0.0001, unpaired two-tailed t-test). (C) Schematic outlining the strategy for generating a topological map that integrates single-telocyte transcriptional profiles with spatial position along the crypt-villus axis. Foxl1Cre; Rosa-mTmG mice were used for tissue dissociation, FACS sorting of GFP+ cells, and smFISH mapping for cluster landmark genes. (D) UMAP projection of single-cell RNA-seq data generated from GFP+ sorted jejunal cells (Foxl1Cre; Rosa-mTmG mice). A total of 7,610 cells from 7 mice were categorized into 18 clusters. (E-F) UMAP plots depict Foxl1 and Gli1 expression profiles across the 18 distinct clusters. (G-J) Immunofluorescence in Foxl1Cre; Rosa-mTmG mouse jejunum sections stained for Cd31 (G) and αSMA (H) reveals sparse GFP labeling in pericytes and myofibroblasts alongside subepithelial telocytes. (I-J) smFISH images of Foxl1Cre; Rosa-mTmG mouse jejunum, hybridized for Foxl1 (white dots), show Foxl1 mRNA in GFP+ (green) pericytes (I) surrounding large vessels (V) beneath the muscularis mucosa, and in myofibroblasts within the villus core (J, asterisk marks GFP-labeled myofibroblast).

To explore telocyte heterogeneity, we performed single cell RNA sequencing (scRNA-seq) on jejunal tissue, leveraging the region’s well characterized epithelial landscape along the villus axis^24^. Mesenchymal cells from the jejunum of seven mice were dissociated into single cell suspensions, and telocytes were enriched by FACS-sorting GFP+ populations from Foxl1Cre; Rosa-mTmG mice. Sequencing was carried out using the 10X Genomics platform (**Figure 2C**). After sorting and sequencing, datasets from all mice were integrated and pooled for downstream analysis.

Although this approach increased the telocyte fraction, the dataset also included non-mesenchymal cell types, including endothelial, epithelial, and immune contamination likely resulted from the physical adherence of telocyte fragments or adherent processes remaining attached to neighboring cells during tissue dissociation, and potentially from GFP protein released by lysed telocytes. This is an inherent limitation when sorting cell types with long, fragile cellular processes, and should be considered when interpreting sorted population purity. In total, this process yielded approximately 4,000 mesenchymal cells for downstream analysis (**Figure 2D**).

Within the jejunal mesenchyme, we identified three major cell types: myofibroblasts, pericytes, and telocytes. Foxl1 and Gli1 transcripts were expressed at generally low levels, characteristic of transcription factors, however, both genes were enriched in mesenchymal populations compared to other cell subtypes (**Figure 2E and F**). These findings emphasize the necessity of the Foxl1Cre reporter model for reliable telocyte identification, as transcript levels alone are insufficient. Sparse GFP labeling was observed in pericytes associated with large vessels under the muscularis mucosae (**Figure 2G**) and in a subset of myofibroblasts at the villus-base and mid-villus region (**Figure 2H**), both demonstrating Foxl1 expression (**Figure 2I and J**).

Unsupervised clustering revealed five distinct clusters (**Figure 2D**). One cluster (cluster 9) showed low ribosomal protein gene expression and a reduced fraction of ribosomal gene-positive cells, suggesting compromised cell viability^25^ (**Supplementary Table S1**), this group was excluded from subsequent analyses.

### Single-cell transcriptomics reveals molecular mechanisms of crypt telocytes in the intestinal stem cell niche

The four telocyte clusters representing viable cells exhibited two distinct levels of PDGFRα expression (**Figure 3A**): two clusters with high expression (clusters 0 and 11) and two with low expression (clusters 2 and 15). Immunostaining of jejunal sections from Foxl1Cre-driven GFP mice confirmed this PDGFRα heterogeneity at the protein level (**Figure 3B**). Telocytes at the villus tip and base displayed high PDGFRα levels, while those in the villus mid and crypt regions had lower levels.

**Figure 3.**
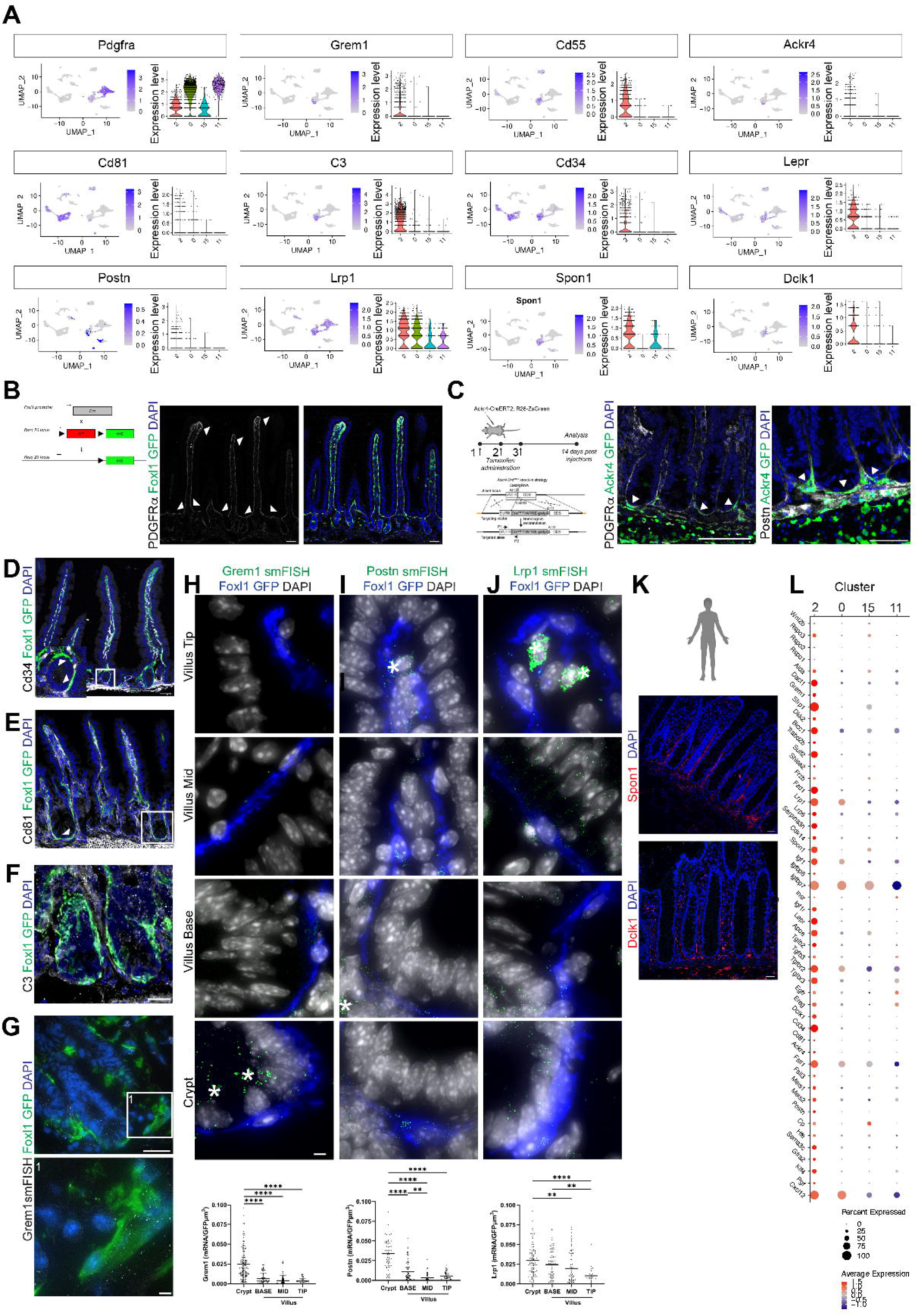
Single-cell transcriptomics reveals molecular mechanisms of crypt telocytes in the intestinal stem cell niche. (A) UMAP and Violin plots display the expression patterns of landmark genes within the 18 clusters identified by single-cell RNA-seq. (B) Immunofluorescence imaging of Foxl1Cre; Rosa-mTmG mouse jejunum sections stained for PDGFRα reveals two subepithelial telocyte subpopulations: high PDGFRα expression at the villus tip and base, and low expression at the villus mid-region and crypt. (C) Schematic representation of the Ackr4CreERT2; Rosa26-ZsGreen mouse model, generated by CRISPR/Cas9 insertion of CreERT2 upstream of the Ackr4 ATG start codon. Immunofluorescence of jejunum sections from this reporter system, stained for GFP and PDGFRα or Postn, demonstrates localization of GFP+ trophocytes below the muscularis mucosa and GFP+PDGFRα+/Postn+ subepithelial telocytes along the crypt region (arrowheads). (D-F) Immunofluorescence analysis in Foxl1Cre; Rosa-mTmG mice stained for GFP and either CD34, Cd81, or C3 shows specific expression within crypt-associated telocytes. (G) smFISH imaging of Foxl1Cre; Rosa-mTmG mouse jejunum hybridized for Grem1 highlights enriched Grem1 transcript patterns in GFP+ crypt telocytes and beneath the muscularis mucosa. (H-J) smFISH images and quantification (n=3 mice/group, <10 crypt-villus axes/bar, ****P<0.0001, unpaired two-tailed t-test) of Foxl1Cre; Rosa-mTmG mouse jejunum hybridized for Grem1, Postn, or Lrp1 (green dots), show specific enrichment of these transcripts in crypt telocytes (blue). Asterisks mark non-specific autofluorescence from immune cells or Paneth cell granules. Scale bar 10μm (K) Immunofluorescence of human colon sections stained for Spon1 or Dclk1 demonstrates subepithelial crypt base localization, correlating with crypt telocytes. (L) Dot plot depicts stem cell niche factor expression in cluster 2 (crypt telocytes), with dot size denoting the percentage of telocytes expressing each gene and color indicating expression levels within the cluster. Scale bar 50μm; smFISH 10μm.

Based on PDGFRα expression, we divided the crypt-villus axis into four segments: crypt, villus base, villus mid and villus tip. We then examined markers associated with the intestinal stem cell niche. Notably, genes such as Grem1, Cd55, Ackr4, Cd81 and C3, previously linked to mesenchymal cells beneath the muscularis mucosae termed trophocytes^11,12^, were enriched in cluster 2 (**Figure 3A**). Additionally, cluster 2 expressed genes previously associated with stem cell niche function, including Cd34^26^ and Lepr^27^. Therefore, we considered the possibility that cluster 2 might represents trophocytes of the muscularis mucosae^11,12^.

To address this issue, we utilized a Ackr4-driven CreERT2 reporter mouse model and examined the distribution of Ackr4+/GFP+ labeled cells in the jejunum following tamoxifen induction. As previously described, Ackr4+/GFP+ cells were broadly distributed beneath the muscularis mucosae (**Figure 3C)**, corresponding to trophocytes^11,12^. Additionally, we detected subepithelial GFP+ signals along the crypts, which co-expressed PDGFRα and Postn, suggesting their identity as telocytes (**Figure 3C** arrowhead). These findings indicate that markers used to identify trophocytes such as Ackr4, can also label crypt telocytes. In contrast, Foxl1 Cre-driven mouse models do not target cells in the muscularis mucosae, highlighting the distinct role of telocytes in the stem cell niche.

Immunostaining for Cd34, Cd81 or the complement component C3 (**Figure 3D-F**), along with smFISH for Grem1, Postn and the apolipoprotein E receptor Lrp1(**Figure 3G-J**) in the Foxl1Cre/ GFP labeled jejunum, identified cluster 2 as telocytes located along the crypt epithelium.

Notably, Grem1 transcripts were predominantly detected in crypt telocytes best captured along the surface region of the crypt, with mRNA distribution extending beneath the muscularis mucosa, resembling the pattern described for trophocytes^11^ (**Figure 3G**). Transcripts lacking GFP colocalization raise the possibility that they may originate from other cell types or from crypt telocytes whose fine cellular processes are not sufficiently visualized. This highlights the need for better cell identification approaches. Use of the Grem1CreERT2 mouse model in combination with the membrane-localized Rosa-mTmG reporter reveals the distribution pattern of telocytes^28^, emphasizing the importance of membrane-based reporter labeling together with smFISH to accurately associate transcripts with specific cell types. Additionally, staining of normal human colon tissue for Spon1 and Dclk1, both enriched in cluster 2, showed localization at the crypt base (**Figure 3K**). This finding further validated the assignment of cluster 2 as crypt telocytes.

Crypt telocytes are enriched for stem cell niche factors as expected, including components of Wnt and R-spondin signaling, as well as Igf, Egf, and Tgfβ, along with diverse trophic factors (**Figure 3L**). Additionally, it expressed markers previously associated with stem cell-supporting functions and colorectal cancer microenvironments. Collectively, these transcriptomic analyses reveal a mechanistic role for crypt telocytes in stem cell niche activity, as demonstrated previously through mouse genetic studies in vivo^7,9,11,26,27,29^.

### Crypt telocyte transcriptomes best resemble stromal cells previously identified as trophocytes

To directly compare the molecular profiles of crypt telocytes and trophocytes, we compared our single-cell data with published trophocyte datasets (GEO: GSM4196131 and GSM4196132). We filtered cells for quality, retaining only those with < 10% mitochondrial transcripts, > 1,500 detected genes, and genes expressed in at least 100 cells, following McCarthy et al., 2020. After merging, scaling, and jointly clustering these datasets using the top 50 principal components (resolution of 0.4), and removing clusters expressing the hematopoietic marker Ptprc (CD45), we identified 14 clusters (**Figure 4A**). Six clusters from the McCarthy dataset showed high similarity to our telocyte clusters.

**Figure 4.**
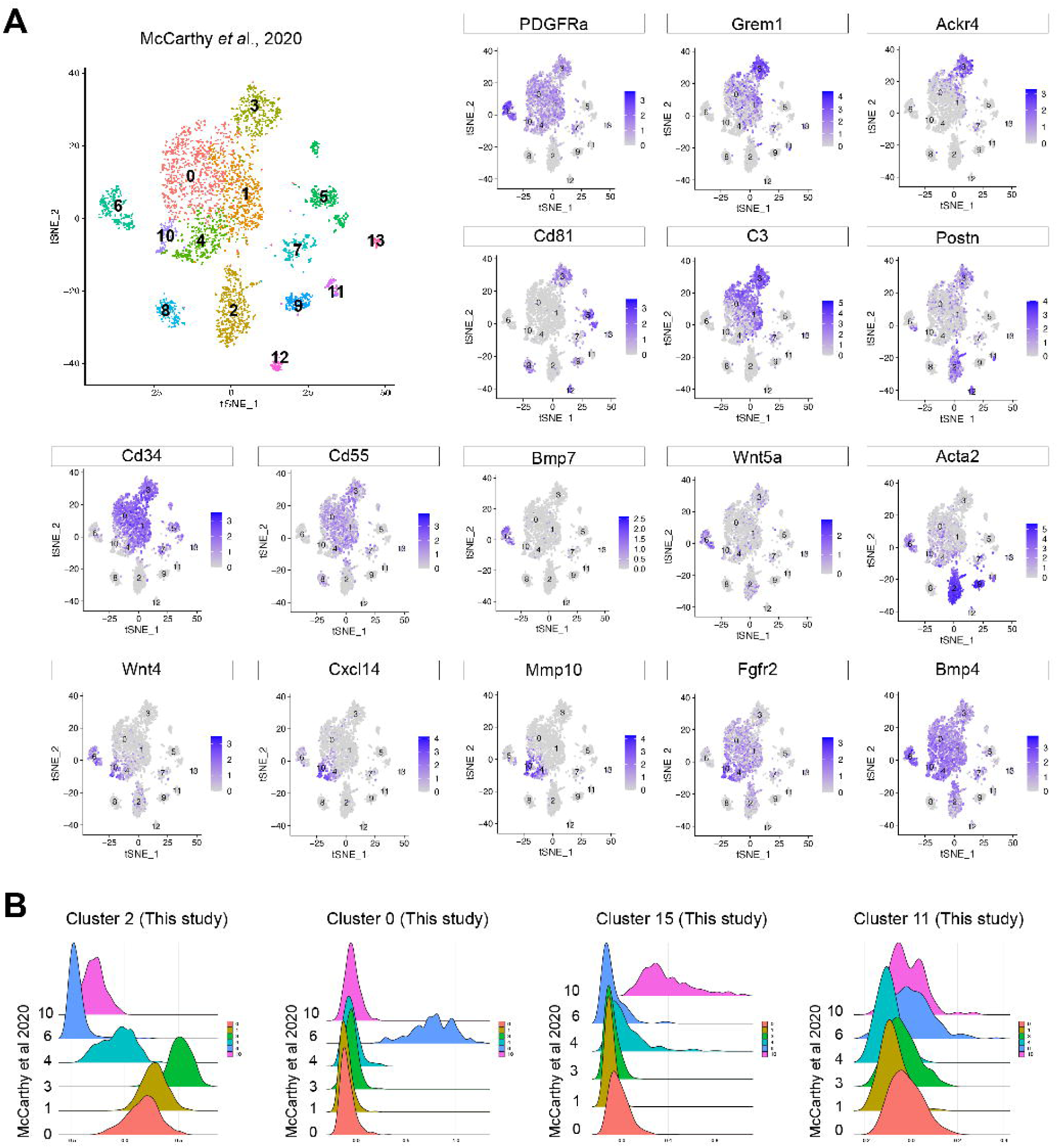
Crypt telocyte transcriptomes best resemble stromal cells previously identified as trophocytes. (A) tSNE projection plots of single-cell RNA-seq data from McCarthy et al., 2020 (GEO: GSM4196131 and GSM4196132), filtered and excluded clusters expressing the hematopoietic marker Ptprc (CD45), published protocol. Fourteen clusters were defined in the McCarthy dataset; six displayed high similarity to the telocyte clusters in this study based on shared marker gene (B) Ridge plot visualizations comparing four telocyte clusters from this study to McCarthy et al., 2020 indicate crypt telocytes (cluster 2) align most closely with three PDGFRα low stromal clusters (clusters 3, 1, and 0) in the McCarthy dataset. Cluster 6 from the McCarthy dataset, previously annotated as telocytes, showed strongest similarity with telocyte cluster 0 from this study, while cluster 15 matched most closely to McCarthy cluster 10 and also showed some similarity to cluster 4.

We calculated module scores for each of the four telocyte clusters using Seurat’s AddModuleScore and visualized these via RidgePlot (**Figure 4B**). The expression patterns of key crypt telocyte markers (Grem1, Ackr4, Cd81, C3, Postn, Cd34 and Cd55; **Figure 4A**), and module scores (**Figure 4B**) matched most closely to three PDGFRa-low stromal clusters (clusters 3, 1 and 0), including both Cd81+ trophocytes (cluster 3) and Cd81-negative stromal cells^12^ (clusters 1 and 0). Cluster 6 in the McCarthy dataset, which was annotated as telocytes, showed the strongest alignment with our telocyte cluster 0, whereas our cluster 15 correspond most closely to McCarthy cluster 10, with some similarity to cluster 4.

We did not identify a counterpart for our telocyte cluster 11, nor did we detect any population expressing Ackr4 and Grem1 clearly distinct from telocytes, which might indicate the absence of trophocytes beneath the muscularis mucosa in the McCarthy dataset. This discrepancy may be attributable to the manual removal of the muscularis mucosa in McCarthy 2020 study, potentially resulting in the loss of the submucosal mesenchymal compartment where trophocyte reside. Notably, Foxl1 was not detected in the filtered dataset, underscoring the need for genetically engineered models and membrane tagged labeling to faithfully identify telocytes.

### Crypt and villus mid telocytes, as well as villus base and villus tip telocytes, are transcriptionally similar despite their spatial separation

Turning our attention to villus telocytes, we observed a distinct enrichment pattern localized to the villus mid region, corresponding to cluster 15 (**Figure 3L, 5A, 5C, D and G**). Notably, cluster 15 exhibited transcriptional similarities to crypt telocytes (cluster 2). In contrast, clusters 0 and 11 displayed an opposing enrichment pattern, characterized by elevated expression of Bmp5 and the non-canonical Wnt5a (**Figure 5B and G**). This complementary expression extended to PDGFRα, the cell adhesion molecule Alcam, Spon1 and the lectin receptor Cd302 (**Figure 3A and 5G**). Specifically, PDGFRα and Alcam were highly expressed in the villus-base and villus-tip (clusters 0 and 11) but were low in the crypt and villus-mid (clusters 2 and 15), whereas Cd302 and Spon1 followed the opposite trend. Immunostaining of jejunal sections from the Foxl1Cre-driven GFP mouse model further supported these findings, revealing strong Cd302 expression in the crypt and villus-mid (**Figure 5H)**, while Alcam was predominantly expressed in the villus base and villus tip, with lower levels in the crypt and villus-mid (**Figure 5I**). Similarly, smFISH confirmed the enrichment of Wnt2b in crypt and villus mid telocytes while Bmp5 was enriched in telocytes in the villus-base and villus-tip (**Figure 5A and B**), Bmp4 showed relatively uniform expression across telocyte populations, with the highest levels at the villus-tip (**Figure 5G)**. These results further highlight the spatial organization of signaling networks along the crypt-villus axis.

**Figure 5.**
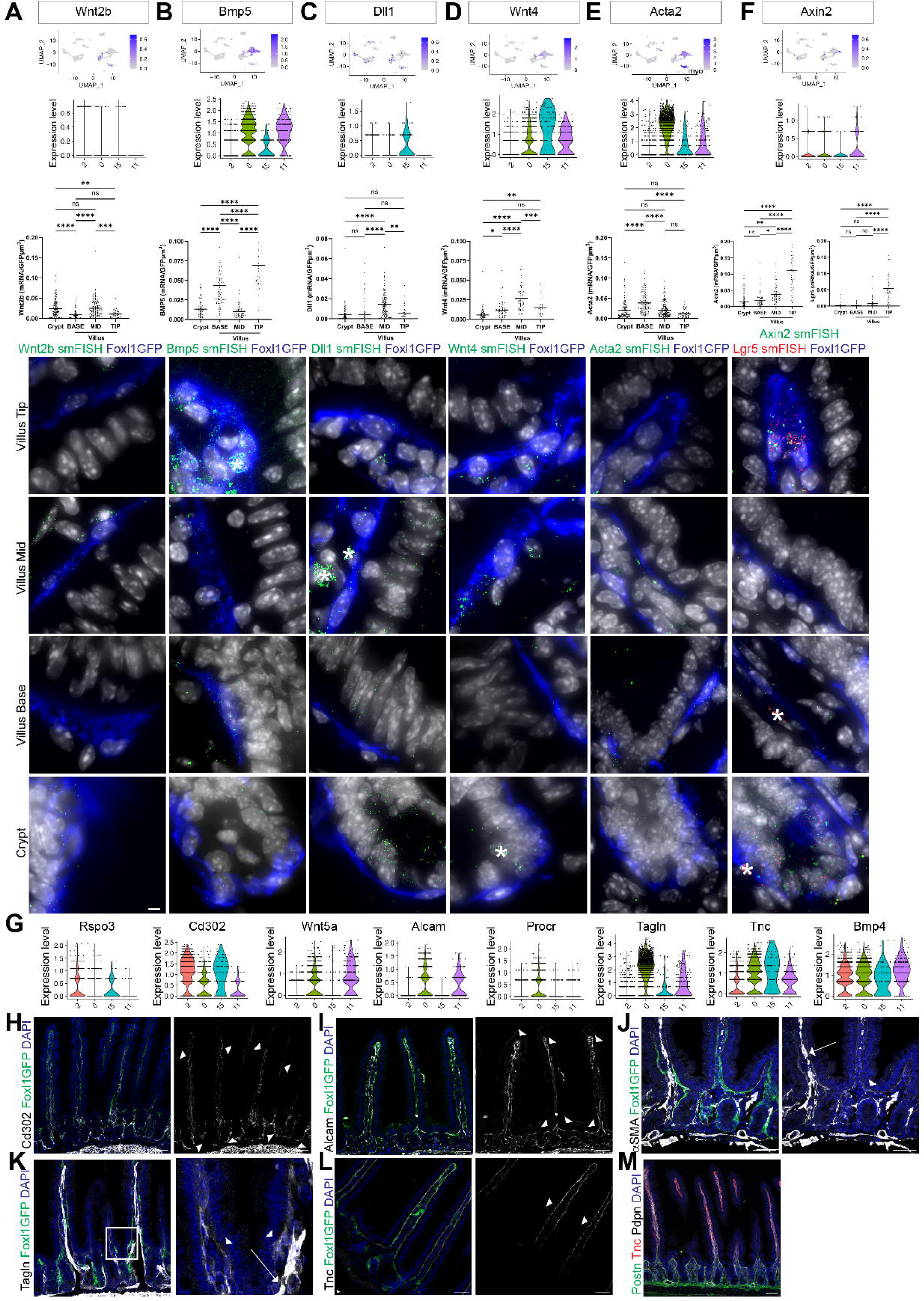
Crypt and villus mid telocytes, as well as villus base and villus tip telocytes, are transcriptionally similar despite their spatial separation. (A-F) UMAP projections, Violin plots, and smFISH quantification/images reveal mRNA concentrations for landmark genes within GFP+ telocytes (blue), located at the crypt, villus base, villus mid, and villus tip. Genes visualized include Wnt2b (A), Bmp5 (B), Dll1 (C), Wnt4 (D), Acta2 (E), Axin2 (green dots) and Lgr5 (red dots) (F). myo-myofibroblasts. Quantification shows significant variation between clusters along the axis (n=3 mice/group, <10 axes/bar, ****P<0.0001, unpaired two-tailed t-test). Asterisks denote non-specific autofluorescence from immune cells or Paneth cell granules. Scale bar 10 μm. (G) Violin plots show landmark gene expression in telocyte clusters, demonstrating complementary patterns: Wnt2b, Rspo3, and Cd302 are enriched in crypt and villus mid telocytes (clusters 2 and 15), while Bmp5, Wnt5a, and Alcam are enriched at the villus base and tip (clusters 0 and 11). (H-M) Immunofluorescence in mouse jejunum sections from Foxl1Cre; Rosa-mTmG (H-L) or C57Bl/6 (M) mice stained for marker combinations. Cd302 is enriched in telocytes along crypt and villus mid regions (H), Alcam is enriched at villus base and tip (I), αSMA and Tagln are enriched at villus base (J, K), Tnc at villus mid (L), and Postn, Tnc, Pdpn at various regions (M). Arrowheads highlight expression in telocytes, arrows indicate expression in myofibroblasts. Scale bar 50μm; smFISH 10μm.

In addition to Wnt2b and Rspo3, cluster 15 (villus mid) was also enriched for the Notch signaling ligand Dll1, the Wnt ligand Wnt4, and the extracellular matrix protein tenascin-C (Tnc) (**Figure 5C, D and G**). smFISH for Dll1 and Wnt4 confirmed the localization of cluster 15 to the villus mid region (**Figure 5C and D**), while immunostaining for Tnc independently validated its highest protein levels in the villus mid (**Figure 5L**).

Interestingly, while Acta2 (αSMA) and Tagln were highly expressed in the myofibroblast population, their expression was also detected in telocytes (**Figure 5E, J and K**). Notably, both markers showed minimal expression in cluster 2, corresponding to crypt telocytes, and the highest expression within cluster 0, although much lower than in myofibroblasts. smFISH for Acta2 (**Figure 5E**) and Immunostaining for αSMA and Tagln (**Figure 5J and K**) support the assignment of cluster 0 to Villus base telocytes. In addition, cluster 0 was also enriched for protein C receptor Procr (**Figure 4G**), while cluster 11 was characterized by high expression of Axin2 (**Figure 5F**). smFISH for Axin2 and Lgr5 showed that telocytes at the villus tip exhibited the highest levels of both markers (**Figure 5F**).

Altogether, our analysis mapped cluster 2 to the crypt, cluster 0 to the villus base, cluster 15 to the villus mid, and cluster 11 to the villus tip, defining a distinct spatial organization of telocyte populations along the crypt-villus axis.

### Differential gene expression analysis suggests distinct functions of telocytes along the crypt-villus axis

Having identified and localized four distinct telocyte populations, we next performed differential gene expression analysis across these four cell clusters (**Figure 6 A-F**). The data revealed distinct patterns of gene activation among the four subtypes as observed by the heatmap of the most variable transcripts (**Figure 6A**). As expected, there is a major distinction between crypt and villus telocytes, but additionally differential gene activation among the telocyte subtypes located in the villus. Specifically, crypt telocytes express multiple components of canonical Wnt signaling, as well as inhibitors of Wnt and Bmp pathways (such as Sfrp1 and Grem1), Hedgehog-interacting protein (Hhip), and elements of the Tgfβ pathway. These cells also exhibit low PDGFRα expression, consistent with their specialized niche function (**Figure 6A-C**). In addition, crypt telocytes showed higher activity of genes involved in extracellular matrix (ECM) remodeling^30^, cell-substrate adhesion, and filament organization. This population also expressed enzymes related to stress resistance, xenobiotic detoxification, and lipoprotein assembly (ApoE) and prostaglandin signaling including Ptger1, Ptger4, Ptgr1 and Ptges (**Supplementary Table 2**).

**Figure 6.**
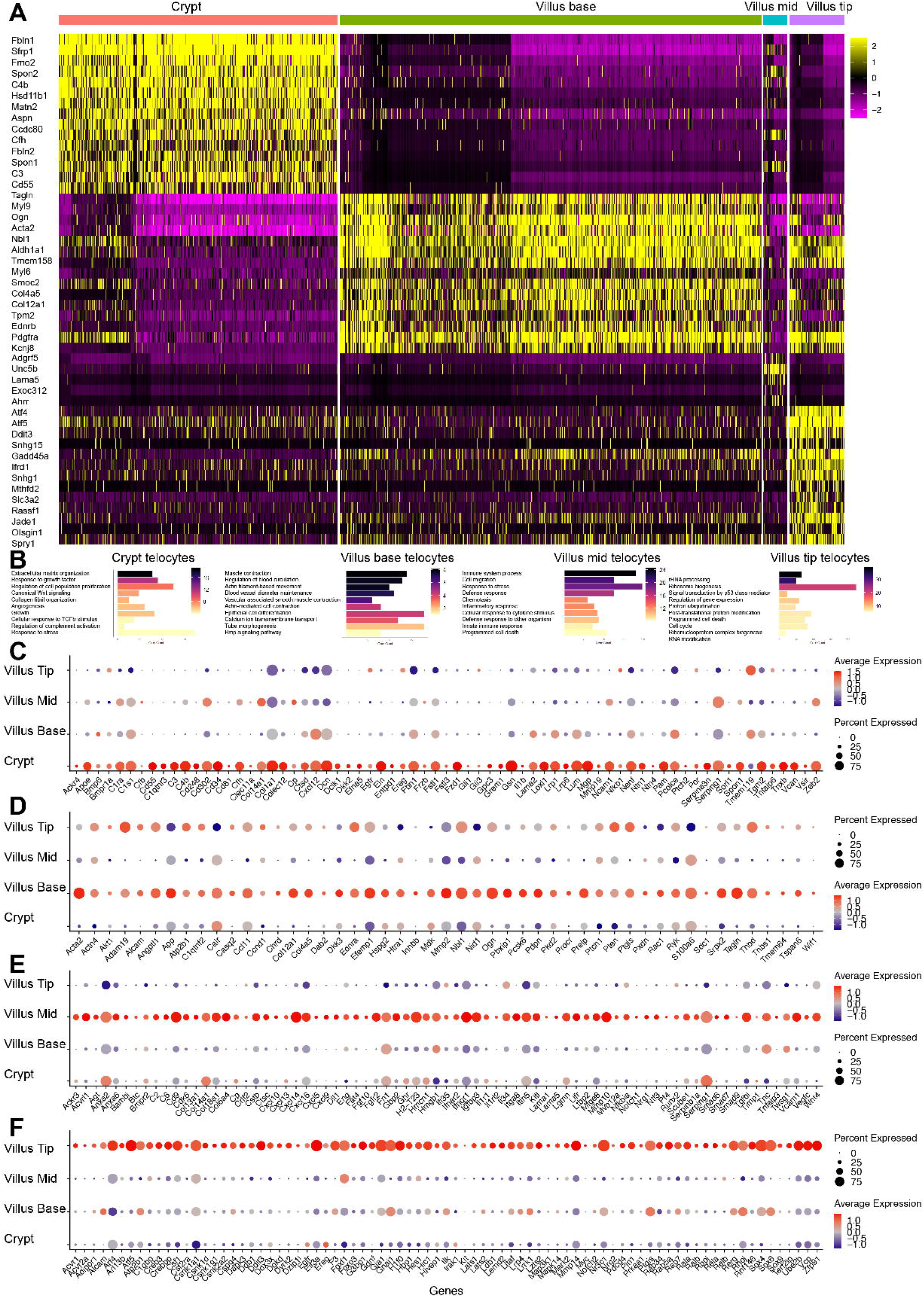
Differential gene expression analysis suggests distinct potential functions of telocytes along the crypt-villus axis. (A) Heatmap showing the main differentially expressed genes in telocyte subtypes located at the crypt, villus base, villus mid and villus tip. Each column represents a single cell, and each row represents a gene. (B) Gene Ontology pathway enrichment analyses of differentially expressed genes in telocyte subtypes at the crypt, villus base, villus mid and villus tip (Biological Process). Scale bars are -Log_10_ (adjusted p-value). (C-F) Dot plots showing the expression of unique genes in telocytes located at the crypt, villus base, villus mid, and villus tip. Each dot represents a gene, with dot size indicating the percentage of telocytes expressing that gene and color reflecting the expression level within each telocyte subtype. This visualization highlights the spatial heterogeneity of gene expression among telocyte populations along the crypt-villus axis.

At the crypt-villus interface, villus base telocytes displayed a transcriptional profile enriched for contractility-related genes, including αSMA, myosin light chain, tropomyosin, and endothelin receptors A and B^31^, suggesting a role in maintaining crypt-villus architecture (**Figure 6A,B and D**). These cells also expressed genes involved in actin cytoskeleton regulation, tube morphogenesis, and blood vessel diameter control, supporting their contribution to tissue structure. Notably, villus base telocytes were enriched for genes involved in retinoic acid metabolism and Bmp signaling (e.g., Aldh1a1, Aldh1a2, Aldh2, Bmp3, Bmp5, Bmp6 and Bmp7), which may help establish the villus-to crypt Bmp gradient that regulates epithelial differentiation.

Villus mid telocytes, though small in number, exhibited a profile indicative of immune surveillance and inflammation, with upregulation of genes involved in chemokine and cytokine signaling, host defense, and adaptive and innate immunity (**Figure 6A, B and E**). This included signatures for interferon responses, Il-6/JAK/STAT3, and Tgfβ signaling, suggesting a key role in modulating inflammatory responses and tissue homeostasis.

Finally, villus tip telocytes exhibited a unique transcriptional landscape characterized by enrichment of genes involved in ribonucleoprotein complexes, ribosome biogenesis, nutrient sensing, and the regulation of autophagy (**Figure 6A, B and F**). These cells also demonstrated high expression of transcription factors including Foxf1, which is closely linked to and co-regulated with Foxl1^32,33^, NFκB and p53, as well as genes linked to the Tnfα, mTOR, and Wnt/β-catenin signaling pathways, suggesting roles in metabolic adaptation and cellular stress responses. Furthermore, villus tip telocytes expressed components of chromatin remodeling factors and polycomb repressive complex components, protein ubiquitination and post-translational modification pointing to a potential role in gene and protein regulation and maintenance of epithelial integrity.

### Spatially Zonated potential functions of intestinal telocytes: growth, matrix, and immune regulation along the crypt-villus axis

Distinct extracellular matrix (ECM) gene expression profiles were observed among telocyte subtypes (**Figure 7A**). Crypt telocytes predominantly expressed genes involved in collagen type I synthesis and filamentous network formation, whereas villus telocytes were enriched for basement membrane gene expression. Immune-related gene also followed a zonated distribution (**Figure 7B**). Villus tip telocytes were enriched for genes mediating pathogen recognition and innate immune activation, while villus mid telocytes mainly expressed chemokine and cytokine signaling genes. Notably, crypt telocytes showed high expression of complement cascade proteins, which promote antibody- and phagocyte-mediated clearance of pathogens and damaged cells. These data suggest that telocytes play a protective immune role for crypt stem.

**Figure 7.**
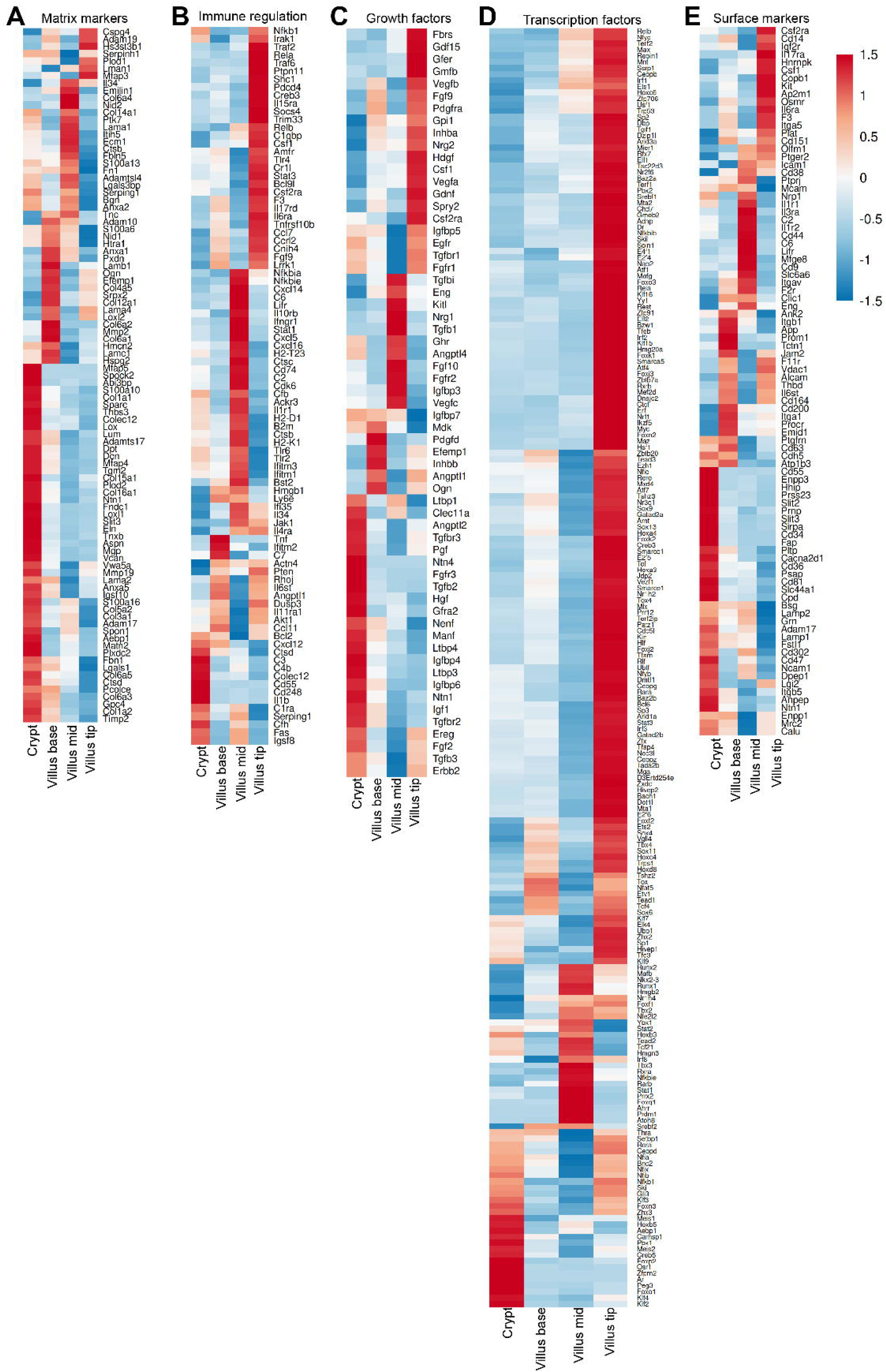
Spatially Zonated potential functions of intestinal telocytes: growth, matrix, and immune regulation along the crypt-villus axis. (A-E) Heatmaps of bulk RNA-sequencing data from single cells illustrate gene expression patterns of the matrix markers (A), immune regulation (B), growth factors (C), transcription factors (D), and surface markers (E) in telocyte subtypes located at the crypt, villus base, villus mid, and villus tip.

Regarding growth factor modulation (**Figure 7C**), crypt and villus-tip telocytes emerged as the main sources of factors such as Egfr, Ereg, Erbb2, Igf-1 and Tgfβ, all of which regulate epithelial growth, proliferation, and differentiation^34–48^. In contrast, villus base and villus mid telocytes primarily produced growth factors that target mesenchymal, endothelial, and immune cells. Villus tip telocytes were uniquely enriched for angiogenic factors Vegfa and Vegfb, which modulate blood vessel fenestration at the villus tip^49^. Conversely, Vegfc, associated with lymphagiogenesis^50,51^, appeared most abundant in villus mid telocytes.

Villus tip telocytes displayed particularly high levels of transcription factors (**Figure 7D**), including Myc-Max, Maf, Mta1, Mta2, Atf1, Etv1 and Bcl6. This population also expressed genes linked to pluripotency, cell cycle control, and chromatin remodeling, indicative of a dynamic, undifferentiated state with potential progenitor-like characteristics.

To aid future translation to human tissues and facilitate telocyte identification, we analyzed the surface marker profiles of telocyte subpopulations (**Figure 7E**). Collectively, these findings reveal pronounced zonation in telocyte functions along the crypt-villus axis, underscoring their specialized roles in ECM remodeling, epithelial morphogenesis, immune regulation, and orchestration of signaling pathways.

### Computational ligand-receptor analysis confirms the major signaling pathways operative between telocytes and epithelial cells

To define the spatial organization of telocyte-epithelial communication along the crypt-villus axis, we analyzed the expression of major signaling pathways from the Wnt and Bmp families (**Figure 8A**), which are central to functional zonation in the intestine^52–56^. Our findings showed that villus tip telocytes predominantly express Bmp ligands. Unexpectedly, these same cells also exhibited strong enrichment of Wnt signaling components, including the β-catenin-Tcf complex, Rnf146, and Tnks, suggesting activation of Wnt signaling through targeted Axin degradation^57^. Villus tip telocytes further expressed casein kinases essential for canonical Wnt signaling in intestinal stem cells^58,59^, as well as key subunits of the β-catenin destruction complex (Gsk3b, Gsk3a, Apc, Ctnnb1, Axin1, and Axin2). The source of Wnt proteins activating this pathway in tip telocytes remins unclear, although villus mid telocytes are promising candidates given their expression of canonical or dual Wnt ligands.

**Figure 8.**
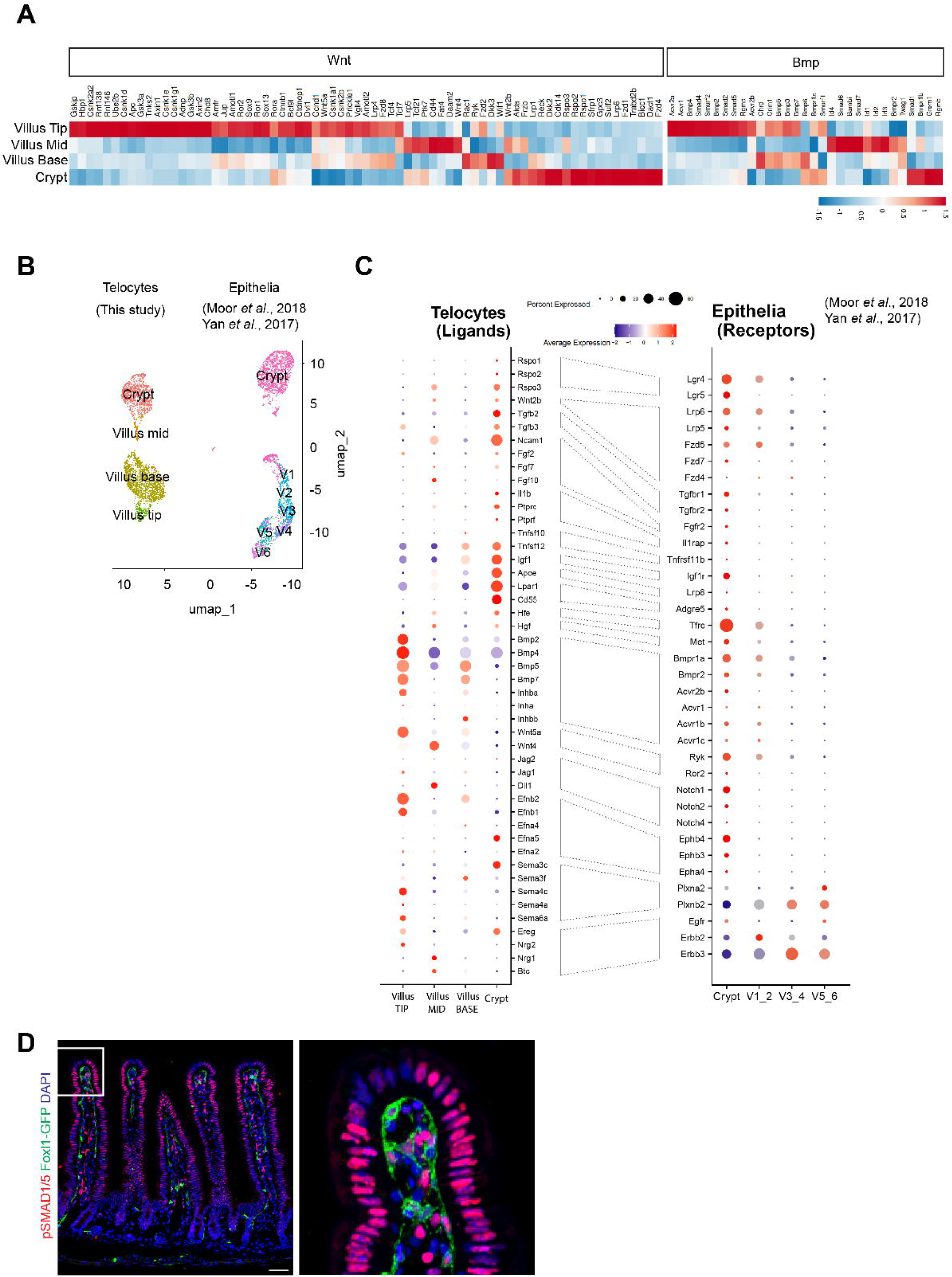
Computational ligand-receptor analysis confirms the major signaling pathways operative between telocytes and epithelial cells. (A) Heatmaps of bulk RNA-sequencing data from single cells illustrate gene expression patterns in the Wnt and Bmp pathways in telocyte subtypes located at the crypt, villus base, villus mid, and villus tip. (B) UMAP plot showing the spatial organization of telocytes and epithelial cells, with each point representing a single cell color-coded by its assigned cluster. V1 to V6 correspond to epithelial regions along the villus axis from villus base to villus tip. (C) Dot plot illustrating predicted ligand-receptor interactions between telocytes (ligands) and epithelia (receptors) across the crypt-villus axis. Dashed lines connect ligands to their corresponding receptors, indicating potential signaling interactions. (D) Immunofluorescence of mouse jejunum sections from Foxl1Cre; Rosa-mTmG stained for GFP and phosphorylated SMAD1/5, indicating Bmp signaling activity, reveals a gradient of Bmp signaling from the villus tip to crypt epithelium, with Bmp activity in GFP+ villus tip telocytes.

BMP signaling, a critical antagonist to epithelial proliferation^60^, showed a distinct gradient: Bmp inhibitors (Grem1, Chrd, Bambi) were mainly localized to crypt, villus base, and villus mid telocytes, with expression lowest in villus tip telocytes. BMP receptor expression followed a similar zonated pattern, peaking at the villus tip, supporting the coordinated regulation of Bmp signaling across telocyte populations. This pattern correlates with the Bmp signaling gradient along the villus-to-crypt axis, highest at the villus tip.

To further resolve ligand-receptor relationships, we used CellphoneDB^61^ to analyze interactions between telocytes and jejunal epithelial cells^24,62^ (**Figure 8B**). Ligands from crypt telocytes correlated closely with the receptors in crypt epithelium, including R-spondins, Wnt, TGFβ, FGF, IL-1, TNF, IGF, HGF, and the iron regulator Hfe. Ligands enriched among villus telocytes primarily had corresponding receptors in crypt epithelium, most notably BMP, non-canonical Wnt, Notch, and Ephrins with Semaphorin and Egfr signaling showing matched zonated activation in villus telocytes and epithelial cells (**Figure 8C**).

To validate these computational predictions, we examined phosphorylation of SMAD1/5, which marks Bmp pathway activation. Although BMP receptors were most abundant in crypt epithelial cells and less so in villus epithelium, phosphorylated SMAD1/5 was highest at the villus tip, confirming that pathway activation is dictated by ligand availability rather than receptor expression. Interestingly, we observed parallel gradients of BMP activation within villus tip telocytes themselves (**Figure 8D**), revealing spatially organized signaling within telocyte populations.

### Ligand-receptor analysis suggests spatial organization of telocyte-to-telocyte signaling along the crypt-villus axis

To investigate how telocyte-to-telocyte communication shapes signaling compartmentalization along the crypt-villus axis, we performed CellPhoneDB analysis to predict ligand-receptor interactions within telocyte populations. In addition to Wnt and Bmp pathways described above, we identified robust TGFβ signaling, critical for intestinal regeneration^63^ and immune regulation^64,65^, particularly enriched in interactions between crypt and villus mid telocytes (**Figure 9**). Crypt telocytes also showed strong connectivity in FGF, IGF, EGF, and Slit-Robo signaling, supporting pro-regenerative functions.

**Figure 9.**
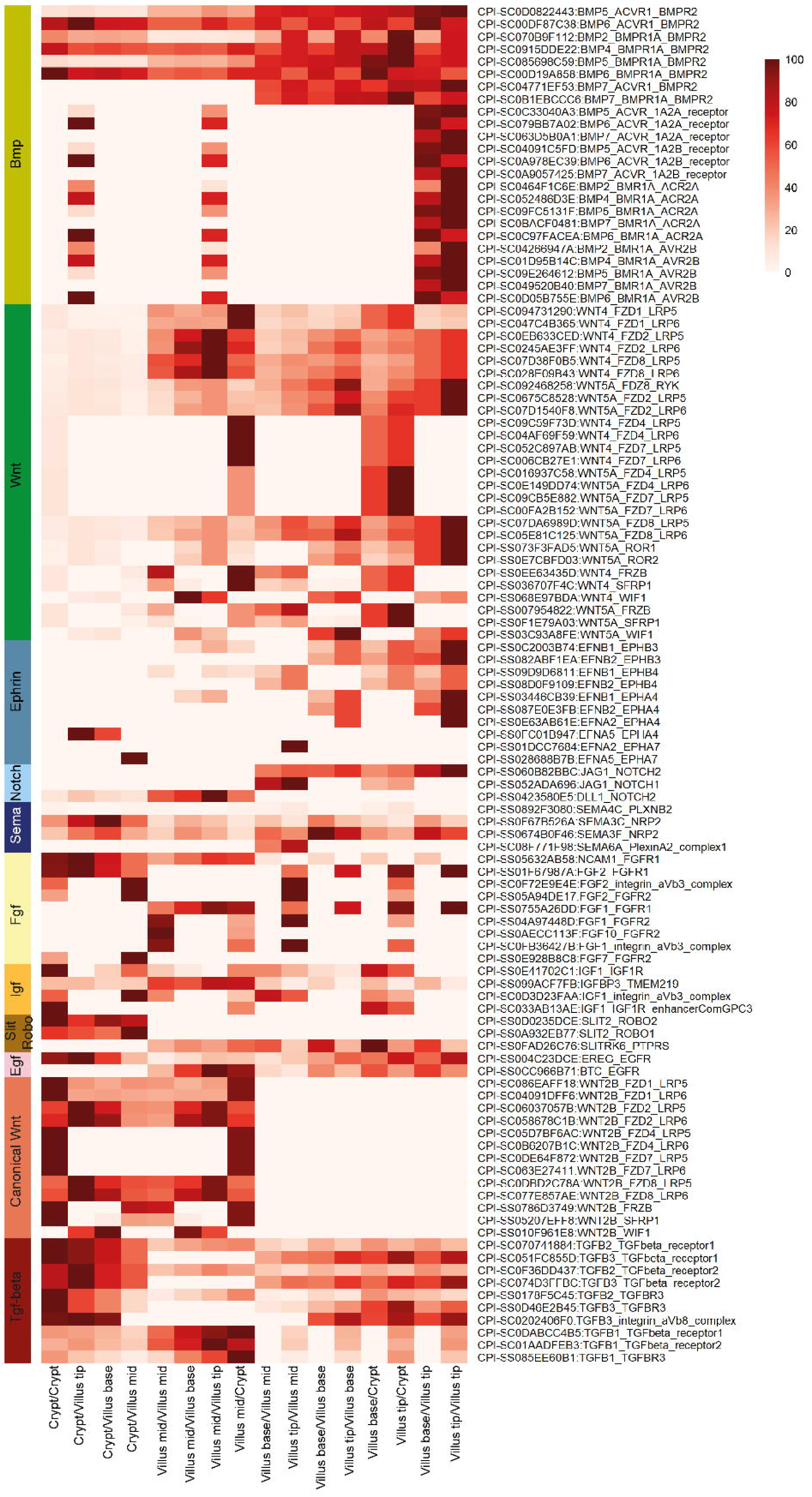
Ligand-receptor analysis suggests that telocyte-to-telocyte signaling is spatially organized along the crypt-villus axis. Heatmap representing the relative activity of signaling pathways across telocyte subtypes. Rows correspond to specific ligand-receptor interactions, identified by unique interaction IDs and ligand-receptor names. Columns represent ligand-receptor interactions specific to each telocyte-to-telocyte interactions. Pathways are color-coded on the left bar for easy identification.

Conversely, non-canonical Wnt, Ephrin, and Notch signaling were biased toward the villus (**Figure 9**), implicating telocytes in Ephrin-mediated epithelial migration^66^ and Notch-driven epithelial differentiation^67,68^. Semaphorin signaling was especially prominent in interactions among villus-base and villus-tip telocytes, both within these subpopulations and between them and crypt telocytes. These findings suggest a previously uncharacterized role for Semaphorin signaling in coordinating telocyte networks along the crypt-villus axis.

Altogether, our results suggest that the spatial architecture of telocyte interactions establishes signaling gradients along the intestinal crypt-villus axis, supporting epithelial homeostasis and compartmentalized function.

## Discussion

A continuous subepithelial fibroblast network, known as the “pericryptal mesenchymal syncytium”, has been recognized since the 1980s based on detailed electron microscopy studies that revealed its three-dimensional architecture^18,69–73^. Despite these foundational morphological insights, the specific functions and cellular identities within this network, especially regarding their regulation of stem cell activity and intestinal regeneration, have remained unclear.

Our study defines the molecular signatures of telocytes along the crypt-villus axis, supporting their role as key coordinators of cellular growth and differentiation in the intestine. Telocytes are distinguished by their unique structural features, most notably long, thin extensions called telopodes, which allow them to form an interconnected three-dimensional meshwork beneath the epithelium^4,74–83^. This network is not homogeneous; rather, telocytes display considerable heterogeneity in gene expression, forming four spatially distinct subpopulations that group into two molecularly similar pairs. Crypt and villus mid telocytes are both enriched for canonical Wnt signaling components, while villus base and villus tip telocytes predominantly express non-canonical Wnt and Bmp ligands. This alternating arrangement suggests that neighboring telocyte subtypes are inter-connected, potentially helping to establish and maintain signaling gradients along the crypt-villus axis. In turn, this organization orchestrates epithelial cell fate decisions and underpins intestinal homeostasis.

Electron microscopy reveals that telocyte extensions form point contacts and electron-dense nanostructures, with adjacent telocytes separated by narrow gaps, suggesting direct molecular interactions and the potential for information transfer across the network^3^. At the molecular level, inferred telocyte-to-telocyte communication via ligand-receptor interactions as reported above further supports the idea of compartmentalized signaling that mirrors the functional zonation of the crypt-villus epithelium.

The transcriptomic similarity between villus mid and crypt telocytes, particularly their shared enrichment for canonical Wnt signaling components likely reflects their parallel roles in supporting proliferative and protective functions within their respective epithelial compartments. While the crypt represents the primary proliferative zone of the intestine, villus mid telocytes may maintain a Wnt-competent state that positions them to support rapid epithelial regeneration during injury or damage. Since Wnt signaling intersects with major inflammatory and regenerative pathways^84–87^, villus mid telocytes likely function as a reserve niche capable of activating regenerative programs when tissue damage demands rapid epithelial repair.

Our ligand-receptor analysis revealed a striking spatial organization: ligands enriched in villus telocytes have their corresponding receptors predominantly localized to crypt epithelium rather than villus epithelium. This apparent spatial segregation does not necessarily indicate direct villus-to-crypt signaling, but rather exemplifies that receptor expression levels alone do not determine where pathway activity occurs. Instead, pathway activity is governed by the combined availability of ligands, pathway inhibitors, and crosstalk with intersecting signaling networks.

BMP signaling provides compelling evidence for this principle. Although BMP receptors are most abundant in crypt epithelial cells, phosphorylated Smad 1/5 the active, downstream mediator of BMP signaling, exhibits a striking gradient with the highest levels at the villus tip (Figure 8D). This striking segregation between receptor distribution and active signaling may be explained by the spatial organization of ligands and inhibitors: BMP ligands are enriched in villus tip telocytes, while BMP inhibitors (Grem1, Chrd, Bambi) are concentrated in crypt, villus base, and villus mid telocytes. Critically, the localization and relative abundance of both ligands and inhibitors, together with inputs from intersecting pathways such as retinoic acid signaling^88–91^, collectively determine where BMP pathway activity is established. Thus, the spatial organization of telocyte-derived signals may create functional signaling gradients that are independent of receptor distribution alone.

Previous attempts to classify the intestinal stroma using scRNA-seq identified telocytes as a uniform population marked by Foxl1 expression. However, the Foxl1 gene is expressed at very low level and therefore is frequently reported as “0” by scRNA-seq, which captures less than 10% of cellular transcripts, even in cells in which the gene is active. This fact and the transcriptomic similarity between telocytes and other stromal cells have made it difficult to distinguish and characterize telocytes without structural validation. By combining scRNA-seq analysis, Foxl1Cre-driven mouse models, and single molecule RNA in-situ techniques, we revealed four telocyte subtypes with distinct potential functions, enriched in specific signaling molecules.

Notably, our data indicate that crypt telocytes, while structurally distinct, are molecularly similar to nearby mesenchymal populations such as Ackr4+ trophocytes, suggesting a possible developmental or functional relationship between these stromal cell types. This raises the intriguing possibility that telocytes may arise from other mesenchymal cells in response to local cues, acquiring their characteristic features to fulfill specialized roles in tissue maintenance and regeneration.

Future experimental approaches that will investigate how telocytes function as a coordinated network, and the specific roles of different telocyte populations in maintaining intestinal homeostasis, will be essential for uncovering new molecular and cellular mechanisms underlying tissue maintenance and regeneration.

## Experimental Methods

### Human samples

The study on human subjects was approved by the Helsinki Institutional Review Board of the Hadassah Medical Center (HMO-0165-19). All patients signed an informed consent.

### Mice

Foxl1-Cre mice^23^ or Foxl1CreERT2 mice^16^ were crossed with the following lines: Rosa-membrane-targeted dimer tomato protein (mT) or membrane targeted green fluorescent protein (mG) (Rosa-mTmG)^92^ (Jackson Laboratories, Bar Harbor, ME #007676) or Rosa-Rainbow^93^. This produced the Foxl1Cre; Rosa-mTmG, Foxl1CreERT2; Rosa-mTmG and Foxl1CreERT2; Rosa-Rainbow.

The Ackr4-CreERT2 mouse model was generated by the Shanghai Model Organisms Center Inc. using CRISPR/Cas9-mediated homologous recombination to insert a CreERT2-WPRE-polyA cassette immediately upstream of the ATG start codon of the Ackr4 gene. Ackr4-CreERT2 mice were then crossed with the Rosa-ZsGreen reporter mice (Jackson Laboratories, Bar Harbor, ME #007906) to produce Ackr4-CreERT2; Rosa-ZsGreen mice.

All animal experiments were approved by the Animal Care and Use Committee of the Hebrew University of Jerusalem.

### Tamoxifen treatment

To induce Foxl1-CreERT2; Rosa-mTmG, Foxl1-CreERT2; Rosa-Rainbow or Ackr4-CreERT2; Rosa-ZsGreen mice, tamoxifen (Sigma-Aldrich #10540-29-1) was dissolved in corn oil at a concentration of 30 mg/ml by shaking overnight at 37 °C. The solution was then administered intraperitoneally at a dose of 150 mg/ kg body weight. Mice were injected once daily for the designated consecutive days and sacrificed for analysis.

### Tissue fixation by trans-cardiac perfusion

Mice were anesthetized by intraperitoneal injection of 7.5% ketamine / 42.5% xylazine (v/v). The thorax and rib cage were then opened to expose the heart. Following this, the right atrium was incised, and a 23G needle connected to a peristaltic pump was inserted into the left ventricle to perfuse pre-cooled (4°C) 4% PFA. Each mouse was perfused with approximately 30 ml fixative. After perfusion, the jejunum or colon were dissected and further fixed with 4% PFA at 4°C overnight.

### Histology

Tissues were embedded in optimum cutting temperature (OCT) compound (Tissue-Tek) for cryosectioning. For the Ackr4CreERT2; Rosa-ZsGreen mouse model, jejunum tissue was embedded in paraffin and antigen retrieval was performed prior to immunostaining.

### Clearing of human and mouse intestines

Intestinal tissues from both humans and mice were fixed in 4% PFA overnight at 4°C and subsequently washed in PBS overnight at 4°C. Tissues were then cleared for 6 hours using X-Clarity^TM^ Electrophoretic Tissue Clearing (ETC) system (LOGOS Biosystems, Annandale, VA), following the manufacturer’s protocol, as previously described^7^.

### Immunofluorescence of cleared whole tissue

Prior staining, cleared intestine was blocked with 10% fetal bovine serum in PBS for 1 hour. Primary and secondary antibodies were diluted in a blocking solution containing 0.3% Triton X-100 and incubated for 48 hours at 4°C each.

Finally, the tissue was mounted *en bloc* on an image slide using 0.5mm thick adhesive silicon isolator mounted in Histodenz solution (88% Histodenz (w/v) in 0.02 M PBS).

### Immunofluorescence staining

For immunofluorescence, tissue was embedded in optimum cutting temperature compound (OCT), and antibodies were diluted in CAS-Block (Invitrogen 008120). Antigen retrieval was performed in a citrate buffer (pH=6) using a pressure cooker (Electron Microscopy Science). All antibodies were diluted in CAS-Block (Invitrogen 008120).

### Antibody list

**Antibody list:**
| Antibody | Catalogue # | Company | Dilution |
| --- | --- | --- | --- |
| Alcam | AF656-SP | Novus Biological | 1:300 |
| C3 | 21337-1-AP | Thermo Fisher | 1:50 |
| CD302 | LS-C801732 | LS Bio | 1:100 |
| CD31 | 557355 | BD Pharmingen | 1:100 |
| CD34 | 14-0341-82 | Thermo Fisher | 1:100 |
| CD45-APC | 103112 | ABCAM | 1:100 |
| CD81-APC | 104909 | Bio Legend | 1:50 |
| E-Cad | 610182 | DB Biosciences | 1:300 |
| EpCAM | ab71916 | ABCAM | 1:500 |
| GFP | AB-ab6673 | ABCAM | 1:500 |
| GFP | NB100-1614 | Novus Biological | 1:250 |
| Lyve1 | MAB2125 | R&D Systems | 1:100 |
| pSMAD1/5 | MAB9516 | Cell Signaling | 1:500 |
| PDGFRα | AF1062 | R&D Systems | 1:100 |
| PDPN-APC | 127418 | Bio Legend | 1:100 |
| Postn | AF2955 | R&D Systems | 1:400 |
| RFP | 600-401-379 | Thermo Fisher | 1:500 |
| TNC | AB19011 | Merck | 1:300 |
| αSMA | ab5694 | ABCAM | 1:250 |
| Tuj1 | BLG-801201 | Bio Legend | 1:500 |
| hPDGFRα | AF-307-NA | R&D Systems | 1:200 |
| Spon1 (F-spon)<br>R4 | Kind gift from<br>Prof. Avihu Klar | References<sup>94,95</sup> | 1:1000 |
| CD45-APC | 103112 | ABCAM | 1:100 |
| CD81-APC | 104909 | Bio Legend | 1:50 |
| Dclk1 | 62257 | Cell Signaling | 1:300 |
| Lyve1 | MAB2125 | R&D Systems | 1:100 |
| pSMAD1/5 | MAB9516 | Cell Signaling | 1:500 |
| H3K9me3 | Ab8898 | ABCAM | 1:100 |

### Quantification of epithelial cells contacted by a single telocye

To quantify the number of epithelial cells contacted by a single telocyte, we analyzed jejunal sections from both Foxl1CreERT2; Rosa-mTmG mice induced with a single tamoxifen injection and Foxl1CreERT2; Rosa-Rainbow mice induced with three tamoxifen injections. For each reporter line, individual telocytes were identified by continuous reporter signal. For crypt region analyses, sections including the basal crypt surface were selected. The intestinal epithelium was labeled by EpCAM immunostaining. Epithelial cells were considered to be in contact with a telocyte if any portion of a reporter-positive telocyte process made contact with an EpCAM+ epithelial cell. The total number of contacted epithelial cells for each telocyte was then counted.

### Single molecule RNA fluorescence in-situ hybridization (smFISH)

We employed Foxl1Cre; Rosa-mTmG mice 6-8 weeks of age. Tissue fixation was performed in 4% PFA. Custom probe libraries, designed using the Stellaris FISH probe designer software, were utilized to hybridize with a desired coding RNA (Bio search Technologies, Inc., Petaluma, CA) and coupled to Cy5, TMR or Alexa 594.

Cryo-sections of 7-micron thickness were prepared for hybridization according to the protocol^96^ with some adaptations. Briefly, tissue sections were digested with proteinase K (10 µg/ml Ambion AM2546) for 10 minutes and washed twice in 2× SSC (Ambion AM9765). Sections were then washed in wash buffer (20% formamide (Ambion AM9342), 2× SSC) for 5 minutes before hybridization with smFISH probes diluted 1:3000 in hybridization buffer (10% dextran sulfate (Sigma D8906), 20% formamide, 1 mg/ml *E. coli* tRNA (Sigma R1753), 2× SSC, 0.02% BSA (Ambion AM2616), 2 mM vanadyl-ribonucleoside complex (NEB S1402S) at 30°C overnight. Following hybridization, sections were washed with wash buffer containing 100 ng/ml DAPI at 30°C for 30 minutes. GFP antibody was diluted in the hybridization buffer, and Alexa 488 secondary antibody was diluted in the GLOX buffer for 20 minutes, followed by DAPI (Sigma-Aldrich, D9542) staining.

Imaging was performed on a Nikon-A1R HD25 microscope with a 100×oil-immersion objective.

### smFISH quantification

The crypt-villus axis of the jejunum was divided into four distinct regions: crypt, villus base (including the crypt-villus junction), villus mid, and villus tip which comprises the upper third of the villus.

Cell segmentation and automatic transcript quantification were performed in the MATLAB program (MATLAB release R2020b, MathWorks) using TransQuant transcript as described previously (Bahar Halpern, 2016).

### Microscopy

Microscopes used in the study:

- Nikon Eclipse Ti2 Confocal System (Nikon, Japan).
- Nikon TI2-LA-FL EPI-FL module for smFISH technology (Nikon, Japan).
- Nikon DS-Fi2 color CCD camera attached to a Nikon Ti automated inverted microscope (Nikon, Japan).

Image processing was performed using Fiji, Imaris v10.0 or NIS Elements software packages.

### Statistics

Statistical and graphical data analyses were conducted using GraphPad Prism 9.5.1. The number of replicates for each experiment and the statistical tests performed are detailed in figure legends.

### Single cell isolation of telocytes

Jejunum tissues were dissected from seven 8-week-old male Foxl1-Cre; Rosa-mTmG mice. Mesenchymal cells were isolated as previously described^7^. GFP+ cells were then isolated by FACS sorting. Viable cells were gated using DAPI exclusion, and doublets were removed based on forward and side scatter profiles. Sorting was performed using Aria III UPG sorter. Epithelial and immune cells were excluded using EpCAM-APC and CD45-APC antibodies.

## Computational Methods

### scRNA-seq analysis

GFP+ sorted cells were loaded onto a Chromium single-cell 3′ v2 chip (10x Genomics) or Chromium Next GEM Single Cell Gene Expression v3.1 (3’) (10x Genomics) and libraries were prepared according to the manufacturer’s protocol. The scRNA-seq libraries were sequenced on the Illumina NextSeq500 sequencing system in a 28-bp and 56-bp as paired-ended.

Seven single-cell datasets (Chromium, 10x Genomics) were filtered, retaining cells with less than 5% mitochondrial gene transcripts, more than 200 but fewer than 6,000 detected genes, and 60,000 UMIs. The resulting dataset included 256, 638, 326, 711, 253, 1204 and 4222 cells per sample. Data were normalized using the Seurat SCTransform (v2) function. Integration was performed using the IntegrateData function in Suerat, generating an integrated object comprising 7,610 cells and 20,292 genes. Principal Component Analysis (PCA) and Uniform Manifold Approximation and Projection (UMAP) were used for dimensionality reduction. Clustering identified 18 clusters at a resolution of 0.4, using the first 50 PCA dimensions and 3,000 integration features. Differentially expressed genes were identified by comparing each telocyte cluster to the other three clusters. The resulting gene lists were analyzed for functional enrichment analysis using the ggplot2 R package to identify unique Gene Ontology (GO) terms. The GO annotation displayed in the figure was selected manually to ensure a diverse representation of biological processes and to minimize repetitive or redundant terms. This approach was taken to provide a clearer and more informative summary of the functional enrichment analysis results.

### Comparing this study dataset to McCarthy et al., 2020

Mouse small intestine whole-mesenchyme single-cell RNA sequencing datasets (GEO samples GSM4196131 and GSM4196132) were processed following the approach of McCarthy et al., 2020. Cells were retained if they showed less than 10% mitochondrial transcript content, contained more than 1,500 detected transcripts, and included genes expressed in ≥100 single cells. Sequencing data were analyzed using Seurat (version 5.2.1) in R (version 4.4.3). After individual filtering, datasets were merged, normalized, log-transformed, and scaled. Clustering was performed using the first 50 principal components at a resolution of 0.4, yielding 20 clusters. Clusters expressing the hematopoietic marker Cd45 (Ptprc) were excluded. This resulted in 14 clusters comprising 3,803 cells, 6 of which were identified as telocyte populations based on marker gene expression. These clusters were further subsetted, and module scores for this study clusters 2, 0, 15, and 11 were calculated using Seurat’s AddModuleScore function. For each telocyte cluster, the top 50 differentially expressed genes (compared with the other 3 telocyte clusters) were identified, and their module scores visualized using Seurat’s RidgePlot function.

### Telocytes-Epithelia ligand-receptor analysis

scRNA-seq datasets of Lgr5-eGFP-negative cells (Chromium, 10x Genomics; accessions GSM2644349 and GSM2644350)^62^ were merged using the Seurat “merge” function. Based on zonation profiles reconstructed as described by Moor et al.,^24^ 1383 cells were assigned to either the Crypt or one of six Villus zones (from villus base to villus tip): Crypt (235 cells), V1 (247), V2 (168), V3 (68), V4 (312), V5 (209), and V6 (144).

The Lgr5-eGFP-positive compartment of these datasets (Chromium, 10x Genomics; accessions GSM2644349 and GSM2644350) was filtered to retain only cells with less than 10% mitochondrial gene transcripts and more than 100 but fewer than 5,000 detected genes. After filtering, 638 and 778 cells from each dataset were added to the 235 Lgr5-eGFP-negative cells annotated as Crypt.

The telocyte clusters (2, 0, 15 and 11) were combined to the Lgr5-eGFP-negative and the two Lgr5-eGFP-positive epithelial cells into a single Seurat object (totaling 5,608 cells). The data were normalized using Seurat SCTransform (v2) and principal component analysis (PCA) and uniform manifold approximation and projection (UMAP) were performed using the first 30 principal components.

Visualization in low-dimensional space (UMAP), as well as Dot plots, Violin plots, and Heatmaps, was performed using Seurat’s built-in visualization functions. Figures were further customized using the ggplot2 R package6 as needed. For bulk RNA-seq analysis, scaled data were retrieved from the data layer of the dot plot objects. Heatmaps of these scaled counts were created using a custom wrapper script for the pheatmap R function. Differentially expressed genes between selected clusters were identified using Seurat’s FindMarkers function, which applies the Wilcoxon rank-sum test by default.Single-cell analysis was performed using Seurat version 5.2.0^97–99^ in R version 4.4.3.

### CellphoneDB predicted ligand-receptor interactions

Relevant cell–cell interactions were classified into two categories: (1) **T|E**, involving only telocyte and epithelial clusters; and (2) **T|T**, among telocyte clusters. Interactions were considered relevant if they included at least, one differentially expressed gene (DEG) and all interacting genes were expressed. For each group, dot plots or heatmaps were generated based on either significant mean expression (mean values shown only if p < 0.05) or interaction scores. Interaction scores (0–100) were calculated using CellPhoneDB v5, which ranks interactions by specificity through a five-step process: filtering low-expressed genes, calculating mean expression, aggregating heteromeric subunits via geometric mean, scaling expression between 0 and 10, and computing the product of scaled values. Notably, interactions can appear relevant but yield low scores due to low expression.

## Data and code availability

Sequencing data have been deposited in the Gene Expression Omnibus (GEO) repository under the accession number GEO: GSE297695

## Acknowledgments

We thank all members of M. Shoshkes laboratory for their helpful and insightful discussions. Special thanks to S. Itzkovitz and his laboratory for the assistance with implementing the smFISH technology, and to K.H. Kaestner and Y. Rinkevich for sharing mouse models. Schematic illustrations were created with BioRender.com

This work was supported by the Israel Science Foundation (grant #1997/19, grant #2908/19 MS-C); The National Natural Science Foundation of China (NSFC) and the Israel Science Foundation (joint grant#3436/20, MS-C, BS), Shanghai Jiao Tong University – The Hebrew University of Jerusalem Joint Seed Funding (HC); HUJI International PhD Talent Scholarship and Isadore Sharp Einstein scholarship for IMRIC outstanding PhD students to A.G; Carole and Andrew Harper Diversity Program to Excellent Palestinian PhD Scholarship to A.J.

## Author contributions

A.G wrote the manuscript, conceived, carried out experiments, analyzed and interpreted the data; N.C carried out single cell isolation and sorting of telocytes for scRNA-seq analysis and performed immunostaining and smFISH experiments; S.E conducted computational analysis of scRNA-seq data; I.P conducted ligand-receptor CellPhoneDB analysis; A.J performed smFISH quantification; M.C conducted gene ontology analysis; M.S collected human samples and performed whole mount tissue clearing and immunfluorescence staining; H.H performed immunofluorescence stainings and processed whole mount images in IMARIS; J.T and H.C performed mouse experiments, immunofluorescence stainings and smFISH experiments; M.A-G and N.S provided human material; M.S.-C and BS designed and supervised the study. M.S.-C wrote the manuscript and directed the study.

---

**Video 1. A single telocyte contacts the crypt base epithelium** Immunofluorescence of a 30μm longitudinal section from the jejunum of a Foxl1CreERT2; Rosa-mTmG mouse, following a single, low-efficiency tamoxifen induction, highlights the structure and spatial distribution of individual telocytes along the crypt region.

**Video 2. Telocytes contact the villus epithelium** Immunofluorescence of a 30μm longitudinal section from the jejunum of a Foxl1CreERT2; Rosa-mTmG mouse, following a single, low-efficiency tamoxifen induction, highlights the structure and spatial distribution of telocytes along the villus region.

**Video 3. Telocytes interconnect along the villus epithelium** Immunofluorescence of a 30μm longitudinal section from the jejunum of a Foxl1CreERT2; Rosa-Rainbow mouse, following three tamoxifen inductions, highlights the inter-connected subepithelial network of telocytes along the villus region.

